# Thermodynamics determines the coupling between growth and byproduct production

**DOI:** 10.1101/2024.01.31.578201

**Authors:** Omid Oftadeh, Vassily Hatzimanikatis

**Affiliations:** Laboratory of Computational Systems Biotechnology, École Polytechnique Fédérale de Lausanne (EPFL), CH 1015 Lausanne, Switzerland

**Keywords:** Thermodynamics, Metabolic Engineering, Mixed Integer Linear Programming, Bilevel Optimization

## Abstract

Genetic manipulation of cells to couple byproduct production and growth rate is important in bioengineering and biotechnology. In this way, we can use growth rate as a selective pressure, where the mutants with higher growth have higher production capacity. Computational methods have been proposed to find knockouts that couple growth and byproduct production. However, none of these methods consider the energetic and thermodynamic feasibility of such knockout strategies. Furthermore, there is no computational study of how variations in metabolite concentrations affect the coupling between growth and byproduct formation. One of the computational methods to find knockouts that couple growth and byproduct formation is OptKnock. OptKnock is a bi-level optimization problem. Here, we integrated thermodynamic constraints into the bilevel formulation of OptKnock to create TOptKnock. We show that the computational efficiency of TOptKnock is comparable to that of OptKnock. TOptKnock can account for the thermodynamic viability of the knockouts and examine how variations in metabolite concentrations affect the coupling. We have shown that the coupling between growth and byproduct formation can change in response to variations in concentrations. Thus, a knockout strategy might be optimal for one intracellular condition but suboptimal for another. If metabolomics data are available, TOptKnock can search for optimal knockout interventions under the given condition. We also envision that the TOptKnock framework will help develop strategies for manipulating metabolite concentrations to couple growth and byproduct formation.

## Introduction

Redesigning and engineering microorganisms to produce valuable biochemicals is an important goal of metabolic engineering. Assuming that microorganisms have evolved to maximize their growth, engineering approaches that genetically couple product formation and growth are more robust. In addition, because biomass formation is accompanied by product formation, microorganisms engineered by such approaches will produce more products over generations by maximizing their growth.

Many computational tools have been developed to find optimal strategies to enhance biochemical production in a host organism. Such methods can suggest strategies for adding heterologous genes^1-3^, performing gene knockouts^4,5^, and suppressing or activating native genes^6^. Many of these methods use constraint-based optimization, where an objective function is optimized, subject to physicochemical constraints.

Bilevel optimization, the optimization of two nested problems, is a popular framework for designing strains with improved product yield. This structure allows searching for optimal interventions with the outer problem while considering that the organism optimizes its physiological objective (usually growth rate) with the inner problem. In this way, we find the interventions that couple product yield with biomass yield. Bilevel problems are nonlinear; however, some can be reformulated into linear problems, such as Linear Programming (LP) or Mixed-Integer Linear Programming (MILP). There are two reformulations for bilevel problems. Two reformulations exist for bilevel problems. One reformulation uses the strong duality^7^, and the other uses Karush-Kuhn-Tucker conditions^8^.

OptKnock is a bilevel method for finding reaction knockout strategies that couple biochemical production and growth rate^7^. OptKnock has been used to design mutant *Escherichia coli* to overproduce malonyl-CoA^9^ and 1,4-butanediol^10^ and mutant *Saccharomyces cerevisiae* to overproduce 2,3-butanediol^11^. Several other methods have been derived from OptKnock with variations in intervention strategy, such as reaction suppression and activation^6^, heterologous reaction addition^12^, and gene deletion^4^.

The OptKnock formulation includes constraints that account for (i) reaction removals, (ii) mass balances, (iii) reaction capacities, (iv) substrate availability, and (v) reaction reversibilities. The latter is imposed as they are in the Genome-scale Metabolic Models (GEMs). The reversibility of the reactions in a GEM is determined based on the standard Gibbs free energies or other available information about the directionality of the reactions^13^. However, assigning reaction directionality in this way can be inaccurate in some cases because the thermodynamic properties of living organisms differ from the standard condition. Thermodynamic-based Flux Balance Analysis (TFA) is a formulation for determining directionalities based on Gibbs free energy under biological conditions^14^.

Here, we integrated thermodynamic constraints into the OptKnock formulation to develop TOptKnock. Then, we recast the nonlinear formulation of TOptKnock as a MILP and used it to find thermodynamically feasible knockout strategies to couple succinate production with growth rate in *E. coli* under anaerobic conditions. Finally, we investigated how the concentration of metabolic cofactors affects the coupling between growth and product secretion. We showed that the performance of a knockout strategy depends on metabolite concentrations and that different strategies may be optimal depending on the abundance of key metabolites. TOptKnock is the first bilevel formulation to consider thermodynamic feasibility, and its development paves the way for incorporating thermodynamics into other bilevel formulations.

## Results

### Computational performance

TOptKnock has additional variables compared to OptKnock, including metabolite concentrations, Gibbs free energies, and reaction directionalities. The number of constraints is also higher due to the thermodynamic constraints. Therefore, solving TOptKnock requires more computational resources. Table 1 lists the number of variables and constraints in the reformulated OptKnock and TOptKnock for iJO1366.

**Table 1:**
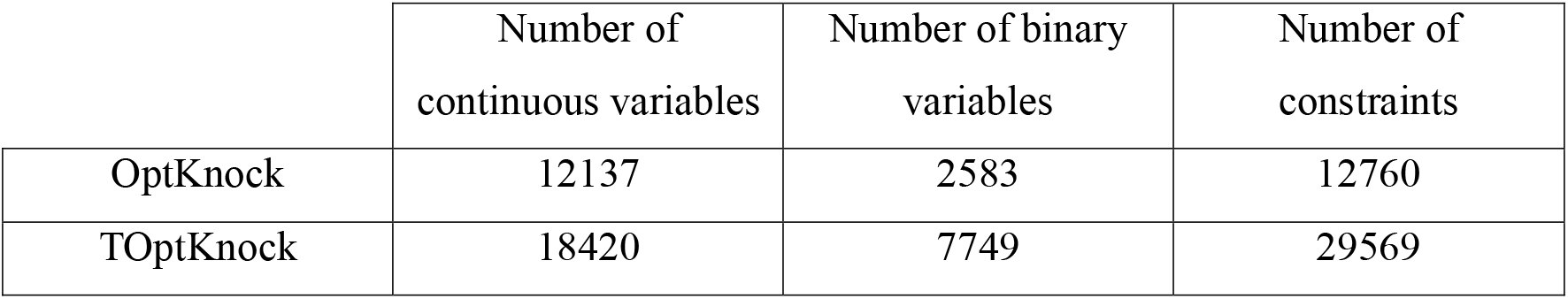
Number of constraints and variables in OptKnock and TOptKnock.

To increase the computational efficiency, we used a numerical trick. We constrained the slack variables for the dual problem (see Methods) to be in the range [0,1]. Based on the formulation, the slack variables are unbounded from above. However, random sampling of the slack variables in the dual problem showed that these variables are usually much smaller than 1. We observed that tightening the bounds on these variables significantly affected the solution time, such that the solution time of TOptKnock was comparable to the original OptKnock (Figure *1*).

**Figure 1:**
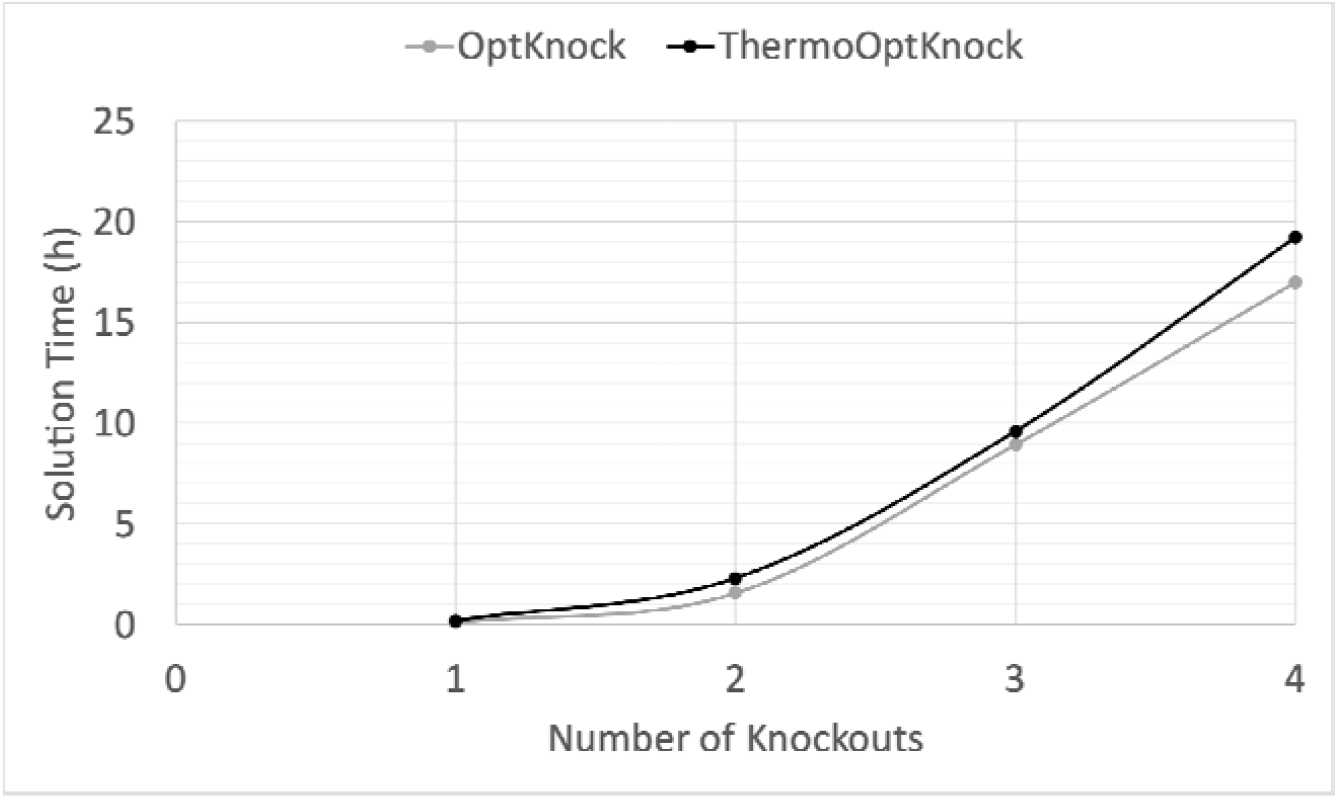
Comparison of the computational performance of OptKnock and TOptKnock. Despite having more constraints and variables, the solution time of TOptKnock was on par with the original OptKnock after tightening the upper bounds of slack variables in the dual problem.

### Using TOptKnock to find knockout strategies

We used TOptKnock to find knockout strategies to couple the biomass and product yield. Such knockout strategies are thermodynamically feasible due to the inclusion of thermodynamic constraints. As a case study, we investigated the overproduction of succinate in *E. coli* under anaerobic conditions. Wild-type *E. coli* scarcely produces succinate at its maximum growth rate (Figure *2*). We generated mutant *E. coli* strains with single, double, and triple knockouts.

**Figure 2:**
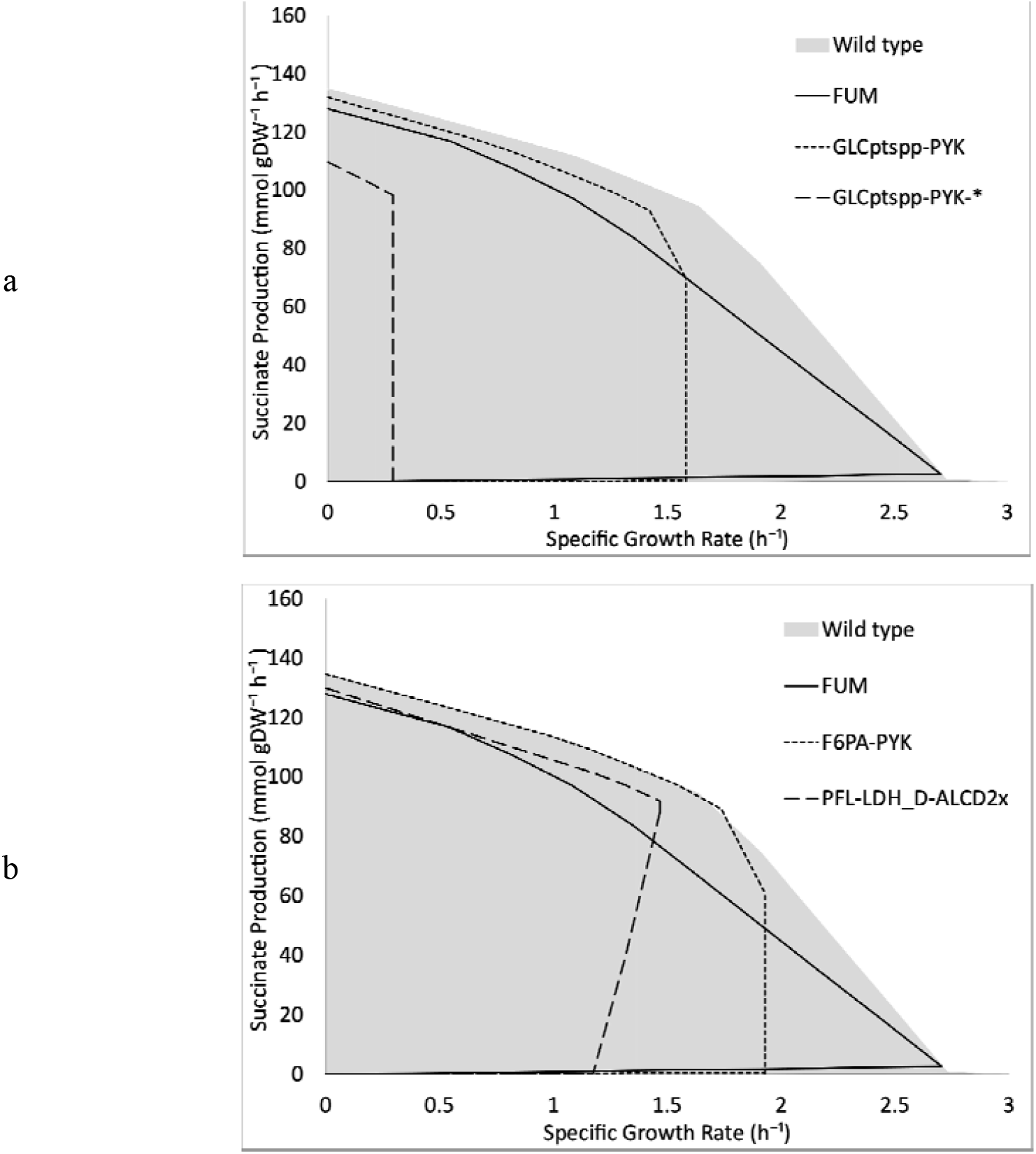
The solution space of different mutant strains. (a) The mutants with the highest succinate production at the maximum growth are compared with the wild-type organism. The only single-knockout mutant with improved succinate production was obtained by removing fumarase (FUM). The strain with removed glucose transport through pyruvate phosphotransferase (GLCptspp) and pyruvate kinase (PYK) had the highest succinate production among the double-knockout mutants. A triple-knockout mutant with removed GLCptspp and PYK (GLCptspp-PYK-*) in addition to either dihydroxyacetone phosphotransferase (DHAPT) or fructose 6-phosphate aldolase (F6PA) had the highest rate of succinate production. (b) The solution space of the mutant strains with the highest OH scores is compared with the wild-type organism. The strain with removed fructose 6-phosphate aldolase (F6PA) and pyruvate kinase (PYK) had the highest OH score (∼11.91) among the double-knockout mutants. The highest OH score (∼131.89) was obtained by the removal of pyruvate formate lyase (PFL), D-lactate dehydrogenase (LDH_D), and alcohol dehydrogenase (ALCD2x).

Removal of the fumarase reaction (FUM), which converts fumarate to L-malate in the TCA cycle, resulted in the only single-knockout mutant with higher succinate production at maximal growth rate (Figure *2*). Increasing the number of reaction knockouts to two and three resulted in more mutant strains with significantly higher succinate yields. Most of these strategies focused on interrupting the conversion of phosphoenolpyruvate (PEP) to pyruvate and diverting PEP towards the reductive branch of the TCA cycle (Figure *2*a).

In all mutant strains, succinate yield was improved at the expense of a reduced biomass yield. We defined the following score to find the knockout mutants with a good trade-off between the biomass and product yields (Figure 3):

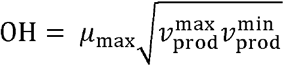

where μ_max_ is the maximum growth rate of the mutant strain. To find 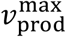 and 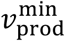, the growth was fixed at its maximum, i.e., μ_max_, and product (succinate) production was maximized and minimized, respectively. We included 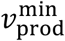 in the definition of OH because higher values of 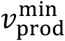 indicate that the coupling between the product and biomass production is forced. In other words, the mutant organism can only increase its growth rate by increasing the production of the product. The OH score for the single knockout mutant with FUM removed was 7.40. The double- and triple-knockout mutants had significantly higher OH scores (Figure *2*b). The highest OH for the double-knockout mutants was 11.91, which was obtained by removing pyruvate kinase (PYK) and fructose-6-phosphate aldolase (F6PA). However, the highest OH, 131.89, was obtained for a triple knockout strategy in which succinate production was tightly coupled to biomass production (Figure *2*b). This strategy prevented the flux from being diverted to the production of alternative byproducts. Removal of pyruvate formate lyase (PFL), D-lactate dehydrogenase (LDH_D), and alcohol dehydrogenase (ALCD2x) blocked the production of formate, lactate, and ethanol, respectively.

**Figure 3:**
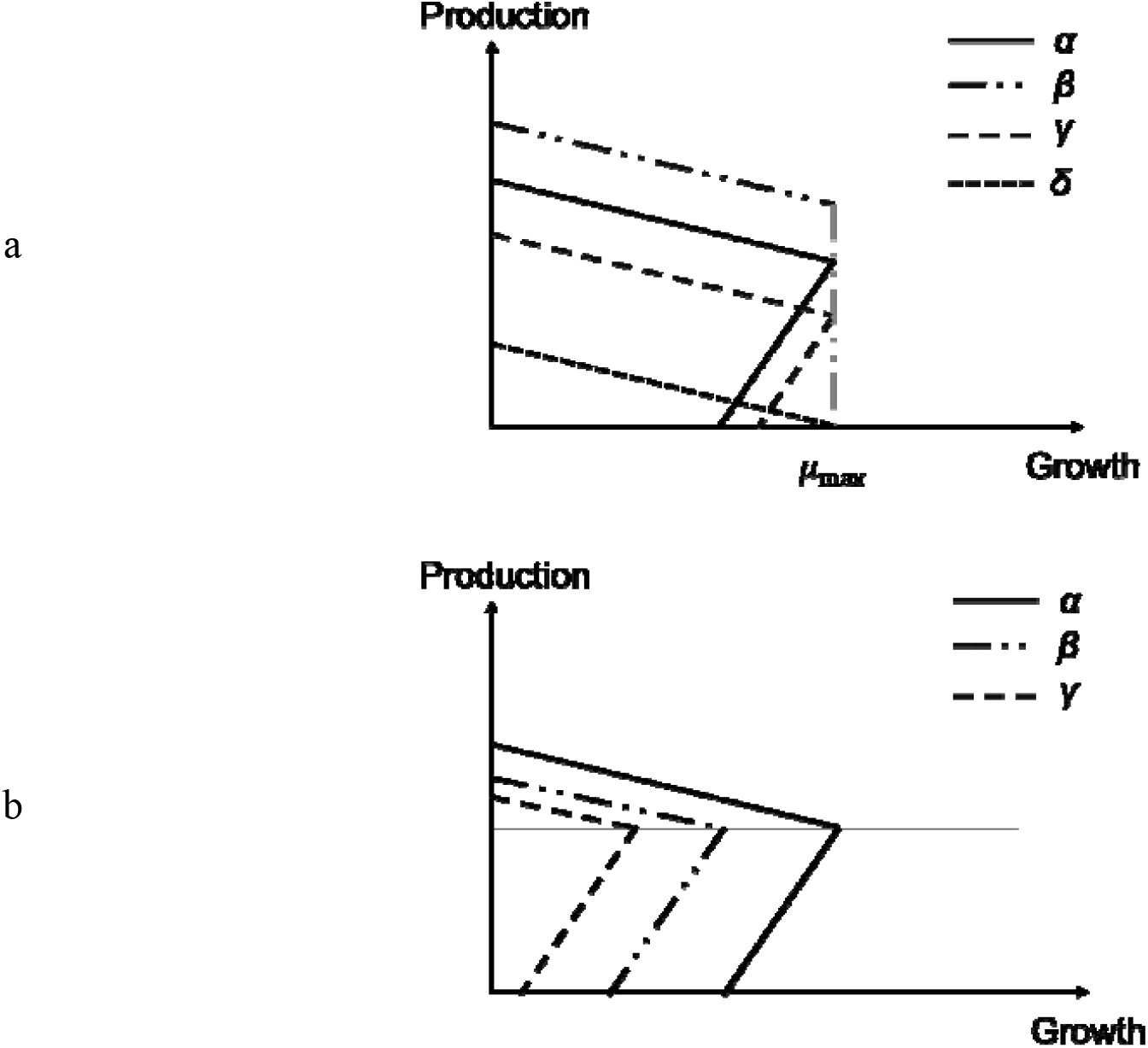
Schematic representation of the OH score. The OH score is defined to rank the solutions, where higher OH scores are preferred. The OH score can increase by increasing the maximum/minimum production rate 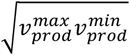 or the maximum growth (μ_*max*_). (a) μ_*max*_ is the same, but the mutant α has the highest OH score due to the higher 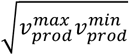. The mutant β shows a higher maximum but lower minimum production rate; in the mutant γ, both minimum and maximum are lower than the mutant α. The mutant δ has the lowest OH score. (b) 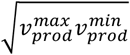 is identical, but the mutant α has the highest OH score due to the higher. Similarly, the OH score for the mutant β is more than the mutant γ.

### The impact of metabolite concentrations on the solution space

The intracellular concentration of key metabolites such as NADH and acetyl-CoA (AcCoA) has been shown to influence succinate production^15,16^. In addition, metabolomic analyses showed that the NAD^+^/NADH and NADP^+^/NADPH ratios differ significantly between aerobic and anaerobic conditions^17-19^. Mainly due to an inactive electron transfer chain, an increased level of NADH is observed under the anaerobic condition^20^. On the other hand, the reduced flux through the oxidative pentose phosphate pathway and the reduced need for protection against superoxide radicals caused a decrease in the level of NADPH^20^. AcCoA/CoA, however, remains almost unchanged^20^.

Such results show how metabolite concentrations can vary in response to changes in metabolism. We performed a sensitivity analysis to evaluate the optimality of different mutants to variations in metabolite concentrations. We generated three mutant strains using the knockout strategies found by TOptKnock to couple succinate production and growth: (i) strain α contains PYK and GLCptspp knockouts, (ii) strain β contains PFL, LDH_D, and ALCD2x knockouts, and (iii) strain γ contains PFL, PYK, and GLCptspp knockouts. These three mutants were the best double knockout (strain *α*), the best triple knockout (strain *β*), and the second-best triple knockout (strain *γ*) mutants. We then constrained the relative concentrations of NAD^+^/NADH, AcCoA/CoA, and NADP^+^/NADPH to be within specified ranges.

In total, we considered 1000 different intracellular conditions, each specified by defining a range for the variation of the key cofactors (Supplementary Table S1). Figure *4* shows the solution space with the highest OH score for each strain and the cofactor ratios for which this solution space is obtained. We assumed that succinate production and growth were tightly coupled if 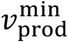 was more than 50% of 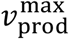 under that condition, and 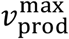 was more than 50% of the maximum succinate production under all conditions. Strains *α, β*, and *γ* showed tight coupling between succinate production and growth in 23, 34, and 52 conditions (Supplementary Table S1), respectively, suggesting that the γ strain is more robust against perturbations in intracellular concentrations.

**Figure 4:**
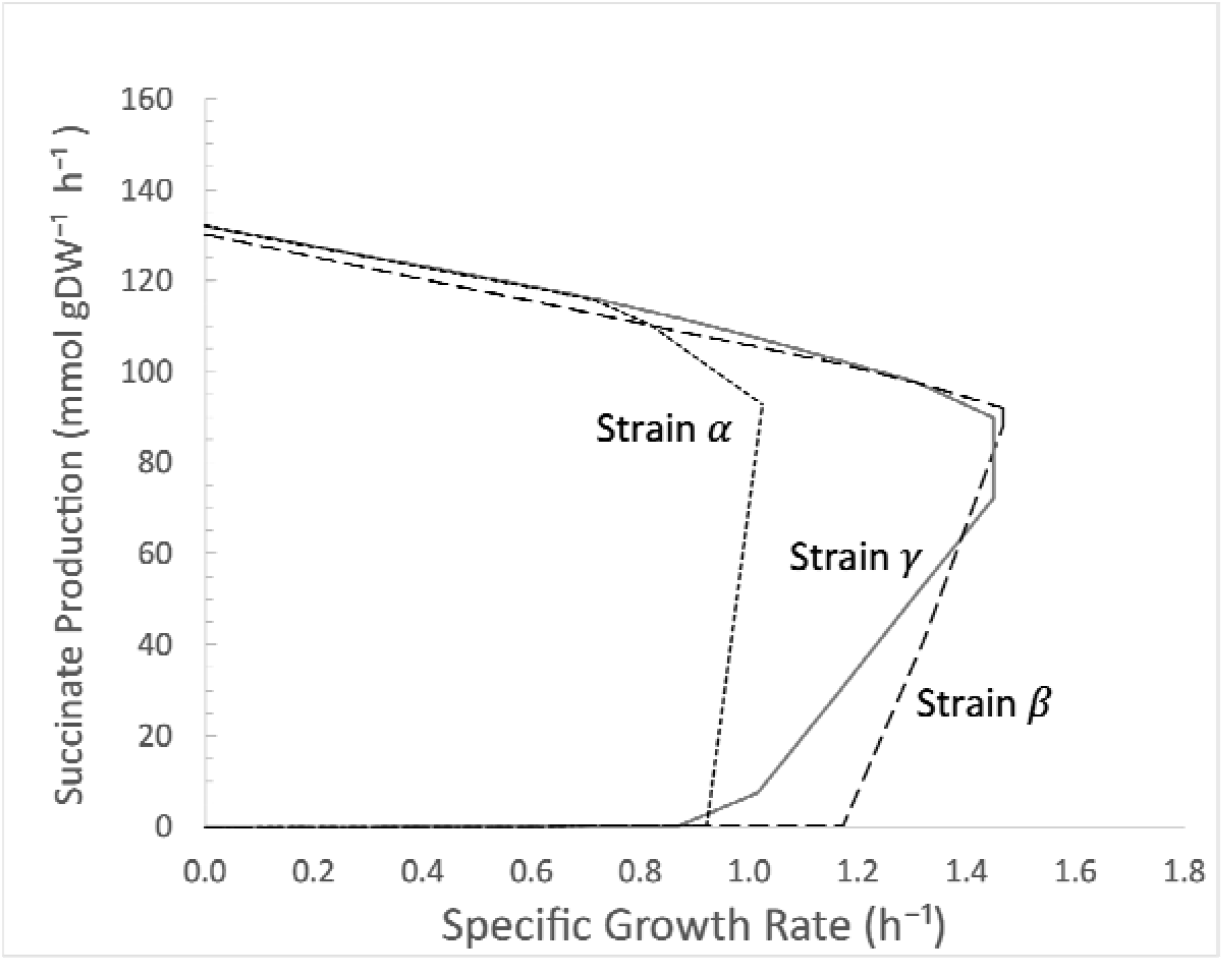
The optimal solution with the highest OH score for each strain. The highest OH score for the *α* strain was obtained when 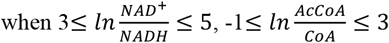, and 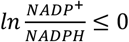. The highest OH score for the *β* strain was obtained when 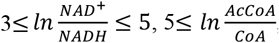 and 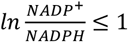. The highest OH score for the *γ* strain was obtained when 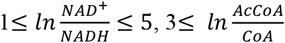, and 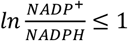.

We specifically explored the solution space for four intracellular conditions (Table *2*) by fixing the growth at different values and minimizing/maximizing the succinate production (Figure *5*). These four conditions were chosen based on our sensitivity analysis (Supplementary Table S1) to demonstrate how an optimal solution under one condition might be suboptimal or non-optimal under another condition. The wild-type organism produces and secretes formate as the main byproduct. However, formate production is blocked in the *β* and *γ* strains because both strains are knocked out for PFL, which in turn causes succinate to be an essential byproduct of growth under condition A. On the other hand, the *α* strain can produce formate; thus, succinate production is not essential under this condition. We performed a variability analysis to find the reaction directions affected after imposing the concentration ranges. Such changes are shown in Supplementary Tables S2-S4.

**Figure 5:**
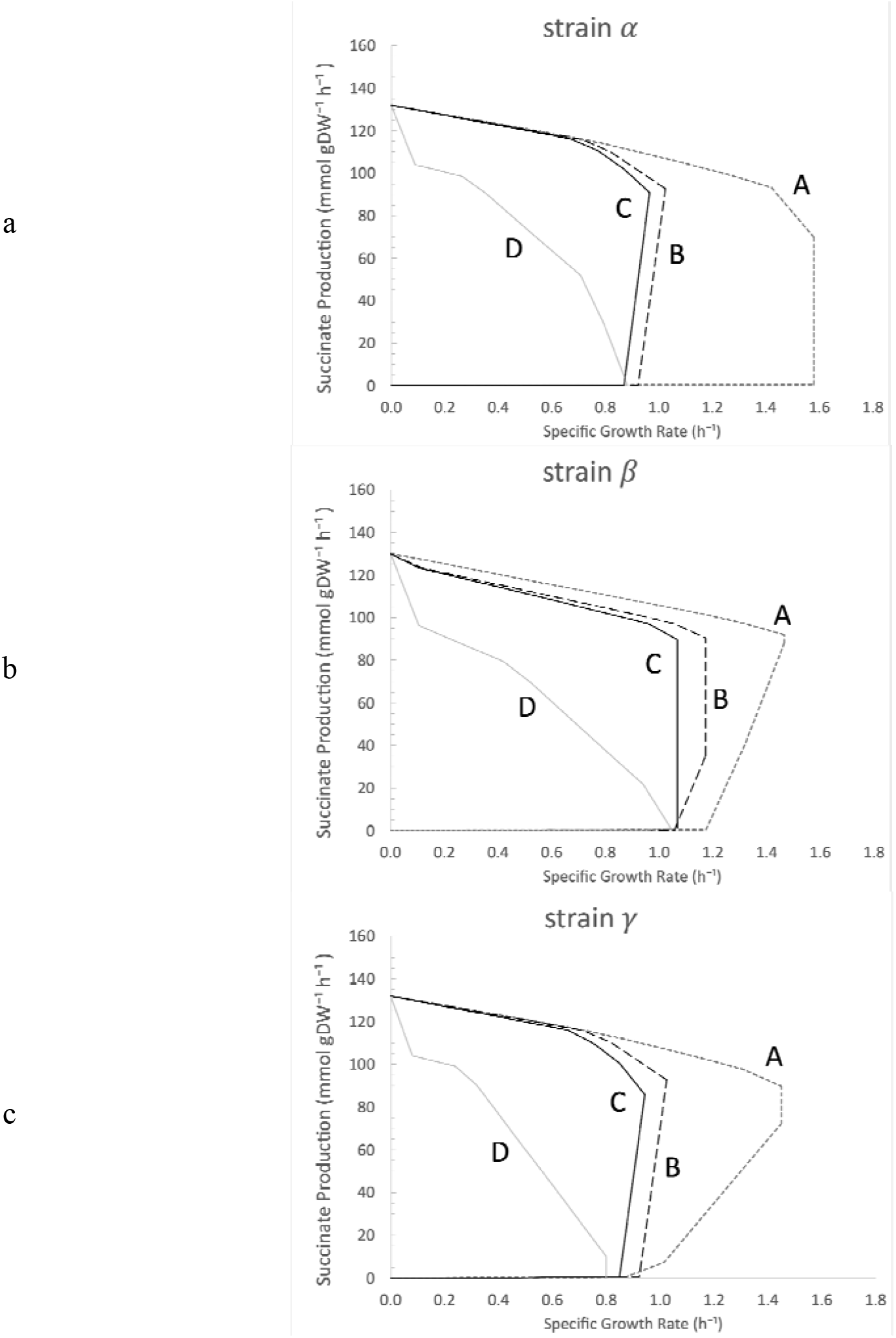
The solution space of the mutant strains under different cellular conditions. (a) The *α* strain was generated by removing GLCptspp and PYK, which interrupted the PEP conversion to pyruvate. Since this strain could produce formate, the succinate production was not tightly coupled to the growth under condition A. In conditions B and D, the AcCoA was less available, adversely impacting the acetate and formate production. Therefore, succinate production was an essential byproduct of the growth in these two conditions. (b) The *β* strain was knocked out for PFL, LDH_D, and ALCD2x. Under conditions C and D, the NAD^+^/NADH was low, which reduced the maximum growth. NADP^+^/NADPH was high under condition C, which diminished the capacity of the cell to produce succinate. The cell could produce succinate under condition D due to the lower NADP^+^/NADPH. However, as the maximum growth rate was lower than the other conditions, other byproducts could be secreted, and succinate production was not essential. (c) The *γ* strain was generated by removing PFL, GLCptspp, and PYK. The removal of PFL blocked the formate production, and succinate was an essential byproduct of growth under conditions A and B. The removal of GLCptspp and PYK interrupted the PEP conversion to pyruvate, which in turn removed the capacity of the cell to produce other byproducts despite the reduced growth under condition D.

**Table 2:** the four intracellular conditions defined by setting bounds on the relative concentrations of key metabolites.

| | $x = \ln \frac{[\text{AcCoA}]}{[\text{CoA}]}$ | $y = \ln \frac{[\text{NAD}^+]}{[\text{NADH}]}$ | $z = \ln \frac{[\text{NADP}^+]}{[\text{NADPH}]}$ |
| --- | --- | --- | --- |
| Condition A | $7 \leq x \leq 9$ | $3 \leq y \leq 5$ | $-3 \leq z \leq -1$ |
| Condition B | $1 \leq x \leq 3$ | $3 \leq y \leq 5$ | $-3 \leq z \leq -1$ |
| Condition C | $1 \leq x \leq 3$ | $1 \leq y \leq 3$ | $-3 \leq z \leq -1$ |
| Condition D | $7 \leq x \leq 9$ | $1 \leq y \leq 3$ | $1 \leq z \leq 3$ |

Condition B, however, forced a much lower availability of AcCoA. As a result, only the reverse direction of the phosphotransacetylase reaction (PTAr) was thermodynamically feasible due to the positive Gibbs free energy. This adversely affected the ability of the cell to produce and secrete acetate and formate, even in the α strain, leaving the cells no choice but to produce succinate. Thus, succinate production was tightly coupled to growth in all mutants in condition B.

Like condition B, condition C featured a low relative concentration of AcCoA, which helped couple growth and succinate production. However, condition C also had lower NAD^+^/NADH than condition B, and it has been previously reported that NADH accumulation negatively affects the growth rate^16,21^. As a result of the reduced growth rate, the coupling between growth and succinate production was not tight in the *β* strain (i.e., 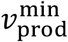 is close to zero). According to the model, the *β* strain can produce L-alanine from pyruvate as an alternative byproduct. On the other hand, strains *β* and *γ* cannot produce L-alanine at maximum growth because the removal of GLCptspp and PYK interrupted the conversion of PEP to pyruvate in these strains. Variability analysis showed that reaction directionalities are identical between conditions B and C except for one reaction (Supplementary Tables S2-S4). The reaction with the changed directionality was 2Fe-2S regeneration (S2FE2SR) in the *α* strain and Octanoate non-lipoylated apo domain ligase (OCTNLL) in the *β* and *γ* strains.

In condition D, the NAD^+^/NADH ratio was the same as in condition C, but the NADP^+^/NADPH ratio was significantly lower. This caused changes in reaction directionalities such that 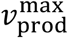 decreased strongly. Thus, succinate production and growth were not coupled in any of the strains under condition D.

We also examined the effect of changing the relative concentrations of each cofactor on the OH score (Figure *6*). We observed no significant differences between different strains in response to the variation in NAD^+^/NADH and NADP^+^/NADPH. The middle ranges of NAD^+^/NADH, i.e., 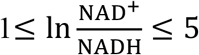, resulted in the highest OH scores. Succinate production was reduced at the higher ratios, while the organisms failed to grow at the lower ratios. On the other hand, we observed a switch-like behavior in response to the change in NADP^+^/NADPH, where at high ratios, i.e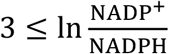, the growth vanished.

**Figure 6:**
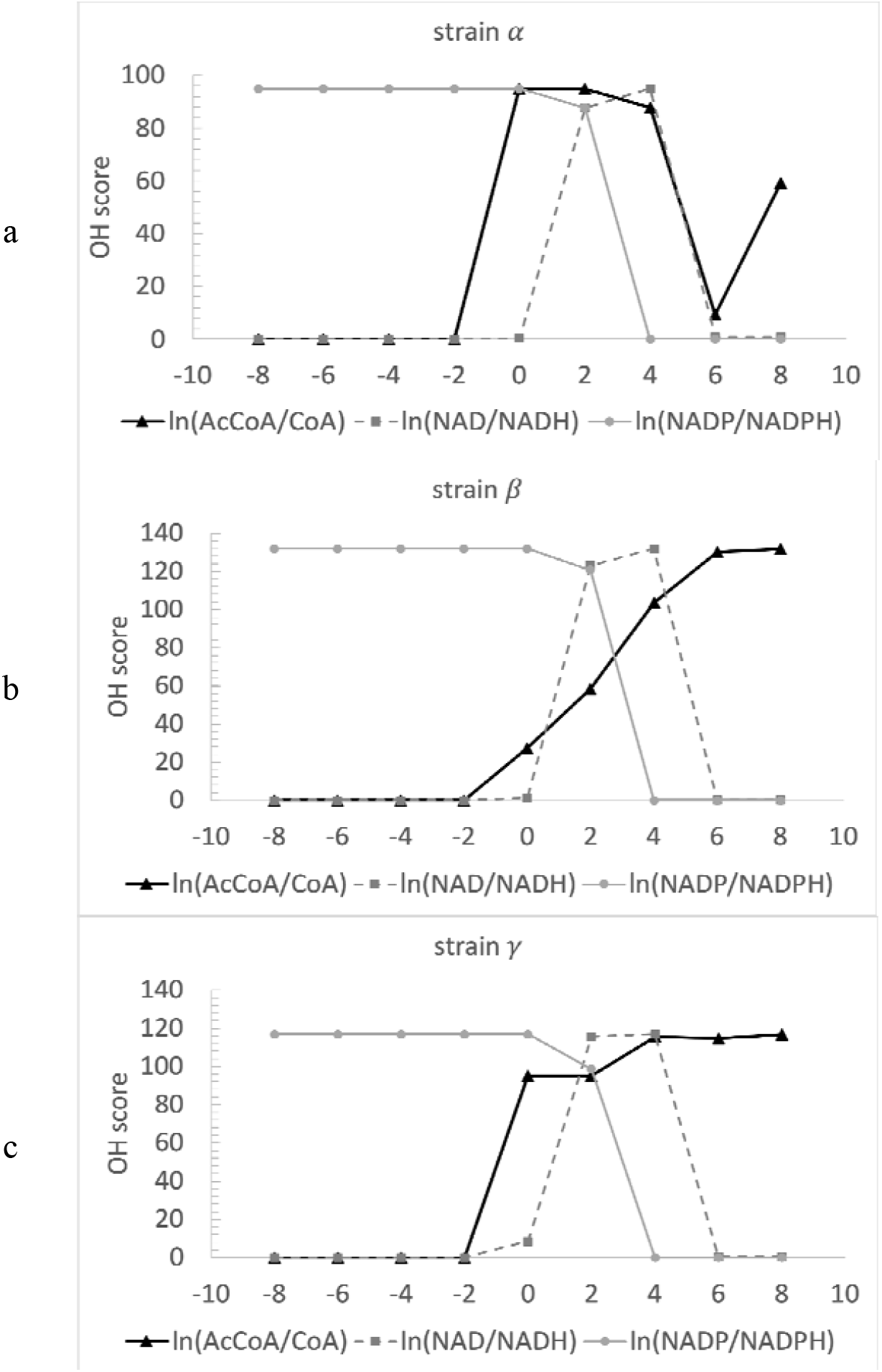
The variation in OH score in response to the changes in AcCoA/CoA, NAD^+^/NADH, and NADP^+^/NADPH ratios. The *α* strain showed a switch-like response to the changes in NADP^+^/NADPH. In response to the variation in NAD^+^/NADH, the OH had a peak for the middle ratios, i.e., 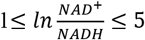. The *α* strain requires lower AcCoA availability to tightly couple the growth and succinate production since this strain is not knocked out for formate production. (b)The *β* strain responded similarly to the *α* strain to the variation in NAD^+^/NADH and NADP^+^/NADPH. Since the *β* strain is knocked out for PFL, this strain cannot produce formate as an alternative byproduct. As a result, even at high AcCoA/CoA ratios, succinate production is tightly coupled to growth. (c) The *γ* strain showed a similar trend to the *β* strain.

Below a certain AcCoA/CoA, i.e., 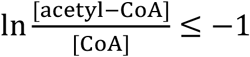, none of the strains could grow. At the higher ratios, however, the strains responded differently to the variations in AcCoA/CoA. Strain *α* had the highest OH scores for the range 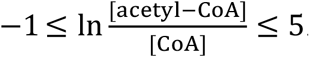. Then, the OH score decreased by increasing the AcCoA/CoA since formate could be produced as the alternative byproduct when AcCoA was sufficiently available. In strains *β* and *γ*, the OH score increased gradually by increasing the AcCoA/CoA due to the increase in the minimum and maximum succinate production.

## Discussion

In this work, we integrated the thermodynamic constraints into the bilevel framework of OptKnock to create a new formulation called TOptKnock. We then recast the bilevel formulation of TOptKnock as a MILP that is solvable using conventional solvers with similar computational resources as the original OptKnock. TOptKnock searches for optimal knockout interventions that (i) are thermodynamically feasible and (ii) couple byproduct production and growth. We have shown that variations in the abundance of key metabolites can significantly affect the coupling between growth and byproduct formation, either by inhibiting growth or affecting the ability to produce the byproduct. The different behavior of the knockout strains under different metabolite concentrations indicates the importance of including thermodynamic constraints in the search for optimal interventions. We observed that a strategy may be optimal under one condition but suboptimal under another. We also observed that some knockout mutants are more robust to perturbations in metabolite concentrations.

In the TOptKnock formulation, the reaction directionalities are determined based on the metabolite concentrations. If metabolomics data are available, TOptKnock can find appropriate interventions for the current cellular state. Furthermore, the TOptKnock framework helps to develop strategies to manipulate metabolite concentrations instead of or in combination with gene knockouts to couple biomass and product yields. Finally, the integration of thermodynamic constraints into OptKnock paves the way for incorporating these constraints into other bilevel methods. This incorporation is of greater importance for methods that involve the addition of novel reactions to a host organism, as the directionality of such reactions in the host is usually not known^1^.

## Methods

### Integration of thermodynamic constraints into the bilevel problem

To determine the directionality of the reactions based on their corrected Gibbs free energy to the biological condition, we integrated thermodynamic constraints to the OptKnock formulation to construct TOptKnock:

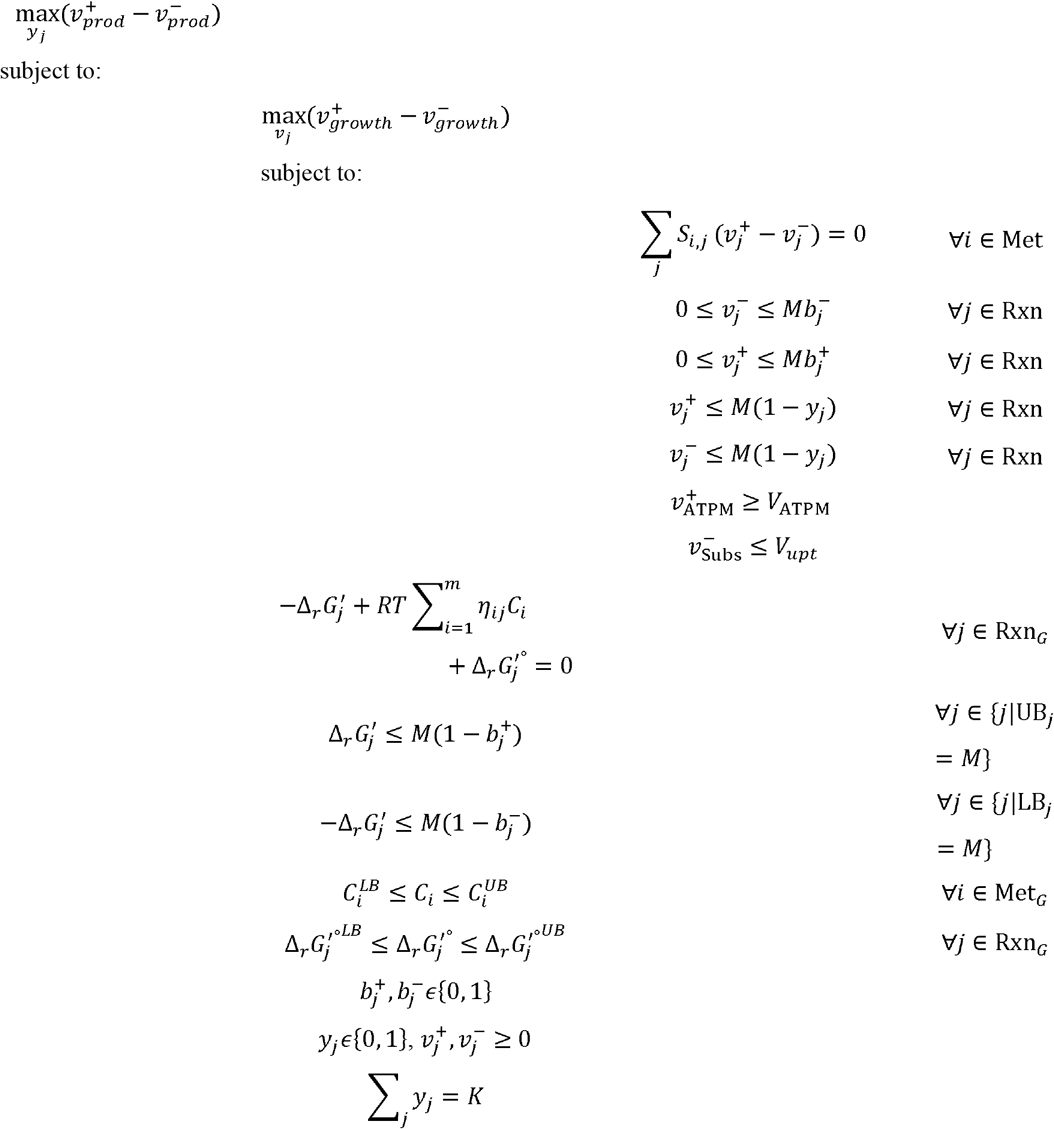

where *C*_*i*_ is the logarithmic concentration of metabolite *i*, 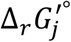 is the Gibbs free energy of reaction *j*. To account for the forward and backward directions, respectively, each flux is represented by two non-negative variables 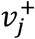 and 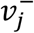. Also, two binary variables 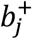 and 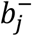 are added to ensure that only one direction is active Met_*G*_ is the set of metabolites with known Gibbs free energy of formation, and Rxn_*G*_ represents the reactions for which thermodynamic constraint is applied.

### Reformulation of TOptKnock

Like OptKnock, we used the strong duality theorem and added the dual constraints and variables of the inner problem to recast ThermoOpKnock as a MILP. However, the reformulation of TOptKnock is not as straightforward as OptKnock since the fluxes are split into forward and backward directions, and additional binary variables are integrated to determine the active directionality. We assumed that 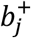 and 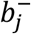 are variables for the outer problem but parameters for the inner problem. A similar assumption was made for reaction knockout variables y_*j*_ in OptKnock^7,22^. The following is the reformulated TOptKnock:

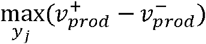

subject to:

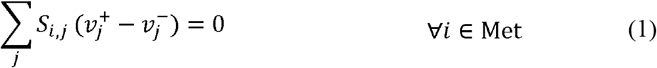

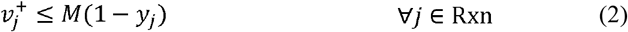

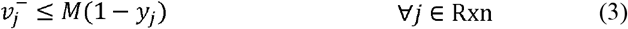

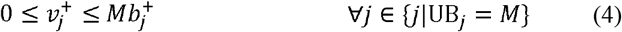

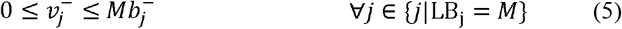

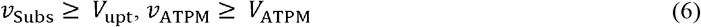

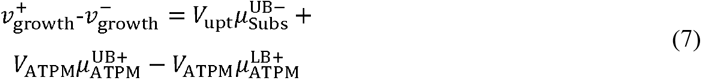

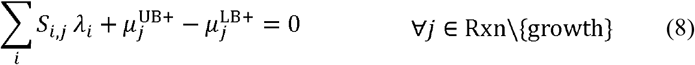

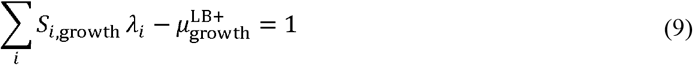

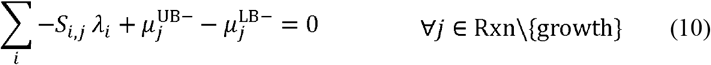

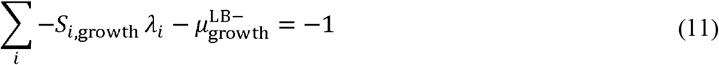

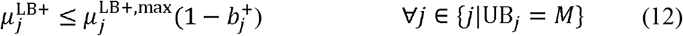

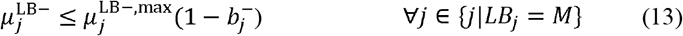

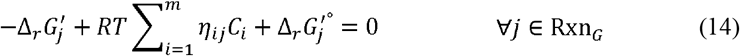

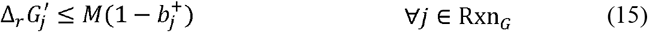

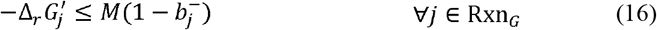

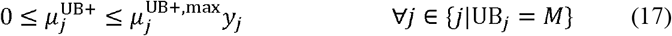

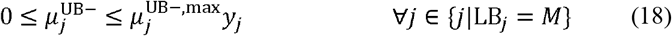

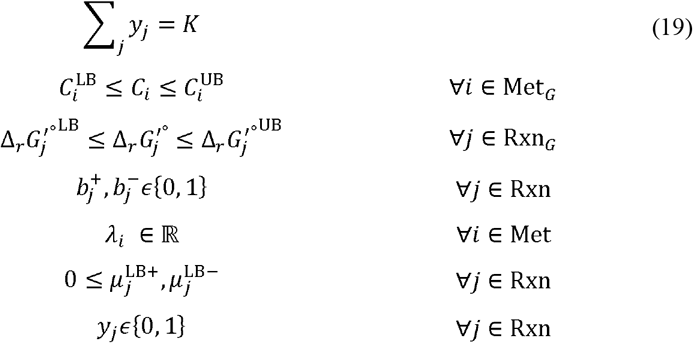

where Equations (1-6 are the primal constraints, Equation (7 enforces the equality of primal and dual objectives, Equations (8-13 are the dual constraints, and Equations (14-19 are the constraints of the outer problem. To increase the computational efficiency, we constrained 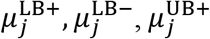 and 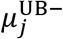 to be in the range [0, 1], which highly reduced the searching space without impacting the optimal solutions.

### Setting up the model for the simulations

The latest version of iJO1366 was obtained from the BiGG database^23^. Uptake of all carbon sources except glucose was blocked. The glucose uptake was constrained to be at most 100 mmol h^-1^ gDW^-1^. The uptake of oxygen was blocked to simulate the anaerobic condition. The lower bound of growth was set to 10% of the maximum growth rate to prevent lethal knockout strategies. All simulations were performed in python 3.7 using the commercial solver CPLEX.

## Acknowledgements

We would like to thank *Dr. Ljubisa Miskovic* for his comments on improving the manuscript. This project was funded by the Swiss National Science Foundation (SNSF): grant 200021_188623, the European Union’s Horizon 2020 research and innovation programme under grant agreement No 814408, and the École Polytechnique Fédérale de Lausanne.

## References

1 Tokic, M. et al. Discovery and evaluation of biosynthetic pathways for the production of five methyl ethyl ketone precursors. ACS synthetic biology 7, 1858–1873 (2018).

2 Kim, J., Reed, J. L. & Maravelias, C. T.Large-scale bi-level strain design approaches and mixed-integer programming solution techniques. PloS one 6, e24162 (2011).

3 Carbonell, P., Parutto, P., Herisson, J., Pandit, S. B. & Faulon, J.-L. XTMS: pathway design in an eXTended metabolic space. Nucleic acids research 42, W389–W394 (2014).

4 Kim, J. & Reed, J. L. OptORF: Optimal metabolic and regulatory perturbations for metabolic engineering of microbial strains. BMC systems biology 4, 1–19 (2010).

5 Patil, K. R., Rocha, I., Förster, J. & Nielsen, J. Evolutionary programming as a platform for in silico metabolic engineering. BMC bioinformatics 6, 1–12 (2005).

6 Pharkya, P. & Maranas, C. D. An optimization framework for identifying reaction activation/inhibition or elimination candidates for overproduction in microbial systems. Metabolic engineering 8, 1–13 (2006).

7 Burgard, A. P., Pharkya, P. & Maranas, C. D. Optknock: a bilevel programming framework for identifying gene knockout strategies for microbial strain optimization. Biotechnology and bioengineering 84, 647–657 (2003).

8 Xu, Z., Zheng, P., Sun, J. & Ma, Y. ReacKnock: identifying reaction deletion strategies for microbial strain optimization based on genome-scale metabolic network. PloS one 8, e72150 (2013).

9 Xu, P., Ranganathan, S., Fowler, Z. L., Maranas, C. D. & Koffas, M. A. Genome-scale metabolic network modeling results in minimal interventions that cooperatively force carbon flux towards malonyl-CoA. Metabolic engineering 13, 578–587 (2011).

10 Yim, H. et al. Metabolic engineering of Escherichia coli for direct production of 1, 4-butanediol. Nature chemical biology 7, 445–452 (2011).

11 Ng, C., Jung, M.-y., Lee, J. & Oh, M.-K. Production of 2, m3-butanediol in Saccharomyces cerevisiae by in silico aided metabolic engineering. Microbial cell factories 11, 1–14 (2012).

12 Pharkya, P., Burgard, A. P. & Maranas, C. D. OptStrain: a computational framework for redesign of microbial production systems. Genome research 14, 2367–2376 (2004).

13 Feist, A. M. et al. A genome[scale metabolic reconstruction for Escherichia coli K[12 MG1655 that accounts for 1260 ORFs and thermodynamic information. Molecular systems biology 3, 121 (2007).

14 Henry, C. S., Broadbelt, L. J. & Hatzimanikatis, V. Thermodynamics-based metabolic flux analysis. Biophysical journal 92, 1792–1805 (2007).

15 Singh, A., Lynch, M. D. & Gill, R. T. Genes restoring redox balance in fermentation- deficient E. coli NZN111. Metabolic engineering 11, 347–354 (2009).

16 Singh, A., Soh, K. C., Hatzimanikatis, V. & Gill, R. T. Manipulating redox and ATP balancing for improved production of succinate in E. coli. Metabolic engineering 13, 76–81 (2011).

17 Partridge, J. D., Scott, C., Tang, Y., Poole, R. K. & Green, J. Escherichia coli transcriptome dynamics during the transition from anaerobic to aerobic conditions. Journal of Biological Chemistry 281, 27806–27815 (2006).

18 Sauer, U., Canonaco, F., Heri, S., Perrenoud, A. & Fischer, E. The soluble and membrane-bound transhydrogenases UdhA and PntAB have divergent functions in NADPH metabolism of Escherichia coli. Journal of Biological Chemistry 279, 6613– 6619 (2004).

19 De Graef, M. R., Alexeeva, S., Snoep, J. L. & Teixeira de Mattos, M. J. The steady- state internal redox state (NADH/NAD) reflects the external redox state and is correlated with catabolic adaptation in Escherichia coli. Journal of bacteriology 181, 2351–2357 (1999).

20 McCloskey, D. et al. A model[driven quantitative metabolomics analysis of aerobic and anaerobic metabolism in E. coli K[12 MG1655 that is biochemically and thermodynamically consistent. Biotechnology and bioengineering 111, 803–815 (2014).

21 Stols, L. & Donnelly, M. I. Production of succinic acid through overexpression of NAD (+)-dependent malic enzyme in an Escherichia coli mutant. Applied and Environmental Microbiology 63, 2695–2701 (1997).

22 Chowdhury, A., Zomorrodi, A. R. & Maranas, C. D. Bilevel optimization techniques in computational strain design. Computers & Chemical Engineering 72, 363–372 (2015).

23 Schellenberger, J., Park, J. O., Conrad, T. M. & Palsson, B. Ø. BiGG: a Biochemical Genetic and Genomic knowledgebase of large scale metabolic reconstructions. BMC bioinformatics 11, 1–10 (2010).

