## Supplementary Tables S2-S4 for "Thermodynamics determines the coupling between growth and byproduct production"

Table 1

Table S2 :Changed reactions directionalities after constraining metabolite concentrations in strain $\alpha$, under different conditions.

| **Reaction IDs** | **Unconstrained metabolite concentrations** | **Constrained metabolite concentrations** |
| --- | --- | --- |
|  | **Condition A** | |
| **2AGPE140tipp** | irreversible in forward | blocked |
| **2AGPE141tipp** | irreversible in forward | blocked |
| **2AGPEAT140** | irreversible in forward | blocked |
| **2AGPEAT141** | irreversible in forward | blocked |
| **ACACT6r** | bidirectional | irreversible in forward |
| **AGPAT140** | irreversible in forward | blocked |
| **AGPAT141** | irreversible in forward | blocked |
| **APG3PAT140** | irreversible in forward | blocked |
| **APG3PAT141** | irreversible in forward | blocked |
| **CLPNH140pp** | irreversible in forward | blocked |
| **CLPNH141pp** | irreversible in forward | blocked |
| **CLPNS140pp** | irreversible in forward | blocked |
| **CLPNS141pp** | irreversible in forward | blocked |
| **CTECOAI6** | bidirectional | irreversible in backward |
| **ECOAH6** | bidirectional | irreversible in forward |
| **FACOAL140t2pp** | irreversible in forward | blocked |
| **FACOAL141t2pp** | irreversible in forward | blocked |
| **G3PAT140** | irreversible in forward | blocked |
| **G3PAT141** | irreversible in forward | blocked |
| **HACD6** | bidirectional | irreversible in forward |
| **LPLIPAL1A140pp** | irreversible in forward | blocked |
| **LPLIPAL1A141pp** | irreversible in forward | blocked |
| **LPLIPAL1E140pp** | irreversible in forward | blocked |
| **LPLIPAL1E141pp** | irreversible in forward | blocked |
| **LPLIPAL1G140pp** | irreversible in forward | blocked |
| **LPLIPAL1G141pp** | irreversible in forward | blocked |
| **PA140abcpp** | irreversible in forward | blocked |
| **PA141abcpp** | irreversible in forward | blocked |
| **LPLIPAL2E140** | irreversible in forward | blocked |
| **LPLIPAL2E141** | irreversible in forward | blocked |
| **PE140abcpp** | irreversible in forward | blocked |
| **PE141abcpp** | irreversible in forward | blocked |
| **PG140abcpp** | irreversible in forward | blocked |
| **PG141abcpp** | irreversible in forward | blocked |
| **PGP140abcpp** | irreversible in forward | blocked |
| **PGP141abcpp** | irreversible in forward | blocked |
| **PGPP140** | irreversible in forward | blocked |
| **PGPP140pp** | irreversible in forward | blocked |
| **PGPP141** | irreversible in forward | blocked |
| **PGPP141pp** | irreversible in forward | blocked |
| **PLIPA2A140pp** | irreversible in forward | blocked |
| **PLIPA2A141pp** | irreversible in forward | blocked |
| **PLIPA2E140pp** | irreversible in forward | blocked |
| **PLIPA2E141pp** | irreversible in forward | blocked |
| **PGSA140** | irreversible in forward | blocked |
| **PGSA141** | irreversible in forward | blocked |
| **PLIPA2G140pp** | irreversible in forward | blocked |
| **PLIPA2G141pp** | irreversible in forward | blocked |
| **PLIPA1E140pp** | irreversible in forward | blocked |
| **PLIPA1E141pp** | irreversible in forward | blocked |
| **PSD140** | irreversible in forward | blocked |
| **PSD141** | irreversible in forward | blocked |
| **PSSA140** | irreversible in forward | blocked |
| **PSSA141** | irreversible in forward | blocked |
|  | **Condition B** | |
| **ACACT6r** | bidirectional | irreversible in backward |
| **ACACT7r** | irreversible in forward | blocked |
| **ACACT8r** | irreversible in backward | blocked |
| **ACOAD7f** | irreversible in backward | blocked |
| **CTECOAI6** | bidirectional | irreversible in forward |
| **CTECOAI8** | irreversible in backward | blocked |
| **ECOAH6** | bidirectional | irreversible in backward |
| **ECOAH7** | irreversible in forward | blocked |
| **ECOAH8** | irreversible in forward | blocked |
| **FACOAE160** | irreversible in forward | blocked |
| **FACOAE181** | irreversible in forward | blocked |
| **HACD6** | bidirectional | irreversible in backward |
| **HACD7** | irreversible in forward | blocked |
| **HACD8** | irreversible in forward | blocked |
| **OCTNLL** | irreversible in forward | blocked |
| **S2FE2SR** | irreversible in forward | blocked |
|  | **Condition C** | |
| **ACACT6r** | bidirectional | irreversible in backward |
| **ACACT7r** | irreversible in forward | blocked |
| **ACACT8r** | irreversible in backward | blocked |
| **ACOAD7f** | irreversible in backward | blocked |
| **CTECOAI6** | bidirectional | irreversible in forward |
| **CTECOAI8** | irreversible in backward | blocked |
| **ECOAH6** | bidirectional | irreversible in backward |
| **ECOAH7** | irreversible in forward | blocked |
| **ECOAH8** | irreversible in forward | blocked |
| **FACOAE160** | irreversible in forward | blocked |
| **FACOAE181** | irreversible in forward | blocked |
| **HACD6** | bidirectional | irreversible in backward |
| **HACD7** | irreversible in forward | blocked |
| **HACD8** | irreversible in forward | blocked |
| **OCTNLL** | irreversible in forward | blocked |
|  | **Condition D** | |
| **2AGPE140tipp** | irreversible in forward | blocked |
| **2AGPE141tipp** | irreversible in forward | blocked |
| **2AGPEAT140** | irreversible in forward | blocked |
| **2AGPEAT141** | irreversible in forward | blocked |
| **ACACT6r** | bidirectional | irreversible in forward |
| **AGPAT140** | irreversible in forward | blocked |
| **AGPAT141** | irreversible in forward | blocked |
| **APG3PAT140** | irreversible in forward | blocked |
| **APG3PAT141** | irreversible in forward | blocked |
| **CLPNH140pp** | irreversible in forward | blocked |
| **CLPNH141pp** | irreversible in forward | blocked |
| **CLPNS140pp** | irreversible in forward | blocked |
| **CLPNS141pp** | irreversible in forward | blocked |
| **CTECOAI6** | bidirectional | irreversible in backward |
| **ECOAH6** | bidirectional | irreversible in forward |
| **FACOAL140t2pp** | irreversible in forward | blocked |
| **FACOAL141t2pp** | irreversible in forward | blocked |
| **G3PAT140** | irreversible in forward | blocked |
| **G3PAT141** | irreversible in forward | blocked |
| **HACD6** | bidirectional | irreversible in forward |
| **LPLIPAL1A140pp** | irreversible in forward | blocked |
| **LPLIPAL1A141pp** | irreversible in forward | blocked |
| **LPLIPAL1E140pp** | irreversible in forward | blocked |
| **LPLIPAL1E141pp** | irreversible in forward | blocked |
| **LPLIPAL1G140pp** | irreversible in forward | blocked |
| **LPLIPAL1G141pp** | irreversible in forward | blocked |
| **PA140abcpp** | irreversible in forward | blocked |
| **PA141abcpp** | irreversible in forward | blocked |
| **LPLIPAL2E140** | irreversible in forward | blocked |
| **LPLIPAL2E141** | irreversible in forward | blocked |
| **OCTNLL** | irreversible in forward | blocked |
| **PE140abcpp** | irreversible in forward | blocked |
| **PE141abcpp** | irreversible in forward | blocked |
| **PG140abcpp** | irreversible in forward | blocked |
| **PG141abcpp** | irreversible in forward | blocked |
| **PGP140abcpp** | irreversible in forward | blocked |
| **PGP141abcpp** | irreversible in forward | blocked |
| **PGPP140** | irreversible in forward | blocked |
| **PGPP140pp** | irreversible in forward | blocked |
| **PGPP141** | irreversible in forward | blocked |
| **PGPP141pp** | irreversible in forward | blocked |
| **PLIPA2A140pp** | irreversible in forward | blocked |
| **PLIPA2A141pp** | irreversible in forward | blocked |
| **PLIPA2E140pp** | irreversible in forward | blocked |
| **PLIPA2E141pp** | irreversible in forward | blocked |
| **PGSA140** | irreversible in forward | blocked |
| **PGSA141** | irreversible in forward | blocked |
| **PLIPA2G140pp** | irreversible in forward | blocked |
| **PLIPA2G141pp** | irreversible in forward | blocked |
| **PLIPA1E140pp** | irreversible in forward | blocked |
| **PLIPA1E141pp** | irreversible in forward | blocked |
| **PSD140** | irreversible in forward | blocked |
| **PSD141** | irreversible in forward | blocked |
| **PSSA140** | irreversible in forward | blocked |
| **PSSA141** | irreversible in forward | blocked |

Table S3 :Changed reactions directionalities after constraining metabolite concentrations in strain $\beta$ under different conditions.

| **Reaction IDs** | **Unconstrained metabolite concentrations** | **Constrained metabolite concentrations** |
| --- | --- | --- |
|  | **Condition A** | |
| **2AGPE140tipp** | irreversible in forward | blocked |
| **2AGPE141tipp** | irreversible in forward | blocked |
| **2AGPEAT140** | irreversible in forward | blocked |
| **2AGPEAT141** | irreversible in forward | blocked |
| **ACACT6r** | bidirectional | irreversible in forward |
| **AGPAT140** | irreversible in forward | blocked |
| **AGPAT141** | irreversible in forward | blocked |
| **APG3PAT140** | irreversible in forward | blocked |
| **APG3PAT141** | irreversible in forward | blocked |
| **CLPNH140pp** | irreversible in forward | blocked |
| **CLPNH141pp** | irreversible in forward | blocked |
| **CLPNS140pp** | irreversible in forward | blocked |
| **CLPNS141pp** | irreversible in forward | blocked |
| **CTECOAI6** | bidirectional | irreversible in backward |
| **ECOAH6** | bidirectional | irreversible in forward |
| **FACOAL140t2pp** | irreversible in forward | blocked |
| **FACOAL141t2pp** | irreversible in forward | blocked |
| **G3PAT140** | irreversible in forward | blocked |
| **G3PAT141** | irreversible in forward | blocked |
| **HACD6** | bidirectional | irreversible in forward |
| **LPLIPAL1A140pp** | irreversible in forward | blocked |
| **LPLIPAL1A141pp** | irreversible in forward | blocked |
| **LPLIPAL1E140pp** | irreversible in forward | blocked |
| **LPLIPAL1E141pp** | irreversible in forward | blocked |
| **LPLIPAL1G140pp** | irreversible in forward | blocked |
| **LPLIPAL1G141pp** | irreversible in forward | blocked |
| **PA140abcpp** | irreversible in forward | blocked |
| **PA141abcpp** | irreversible in forward | blocked |
| **LPLIPAL2E140** | irreversible in forward | blocked |
| **LPLIPAL2E141** | irreversible in forward | blocked |
| **PE140abcpp** | irreversible in forward | blocked |
| **PE141abcpp** | irreversible in forward | blocked |
| **PG140abcpp** | irreversible in forward | blocked |
| **PG141abcpp** | irreversible in forward | blocked |
| **PGP140abcpp** | irreversible in forward | blocked |
| **PGP141abcpp** | irreversible in forward | blocked |
| **PGPP140** | irreversible in forward | blocked |
| **PGPP140pp** | irreversible in forward | blocked |
| **PGPP141** | irreversible in forward | blocked |
| **PGPP141pp** | irreversible in forward | blocked |
| **PLIPA2A140pp** | irreversible in forward | blocked |
| **PLIPA2A141pp** | irreversible in forward | blocked |
| **PLIPA2E140pp** | irreversible in forward | blocked |
| **PLIPA2E141pp** | irreversible in forward | blocked |
| **PGSA140** | irreversible in forward | blocked |
| **PGSA141** | irreversible in forward | blocked |
| **PLIPA2G140pp** | irreversible in forward | blocked |
| **PLIPA2G141pp** | irreversible in forward | blocked |
| **PLIPA1E140pp** | irreversible in forward | blocked |
| **PLIPA1E141pp** | irreversible in forward | blocked |
| **S2FE2SR** | blocked | irreversible in forward |
| **PSD140** | irreversible in forward | blocked |
| **PSD141** | irreversible in forward | blocked |
| **PSSA140** | irreversible in forward | blocked |
| **PSSA141** | irreversible in forward | blocked |
|  | **Condition B** | |
| **ACACT6r** | bidirectional | irreversible in backward |
| **ACACT7r** | irreversible in forward | blocked |
| **ACACT8r** | irreversible in backward | blocked |
| **ACOAD7f** | irreversible in backward | blocked |
| **CTECOAI6** | bidirectional | irreversible in forward |
| **CTECOAI8** | irreversible in backward | blocked |
| **ECOAH6** | bidirectional | irreversible in backward |
| **ECOAH7** | irreversible in forward | blocked |
| **ECOAH8** | irreversible in forward | blocked |
| **FACOAE160** | irreversible in forward | blocked |
| **FACOAE181** | irreversible in forward | blocked |
| **HACD6** | bidirectional | irreversible in backward |
| **HACD7** | irreversible in forward | blocked |
| **HACD8** | irreversible in forward | blocked |
| **OCTNLL** | irreversible in forward | blocked |
|  | **Condition C** | |
| **ACACT6r** | bidirectional | irreversible in backward |
| **ACACT7r** | irreversible in forward | blocked |
| **ACACT8r** | irreversible in backward | blocked |
| **ACOAD7f** | irreversible in backward | blocked |
| **CTECOAI6** | bidirectional | irreversible in forward |
| **CTECOAI8** | irreversible in backward | blocked |
| **ECOAH6** | bidirectional | irreversible in backward |
| **ECOAH7** | irreversible in forward | blocked |
| **ECOAH8** | irreversible in forward | blocked |
| **FACOAE160** | irreversible in forward | blocked |
| **FACOAE181** | irreversible in forward | blocked |
| **HACD6** | bidirectional | irreversible in backward |
| **HACD7** | irreversible in forward | blocked |
| **HACD8** | irreversible in forward | blocked |
|  | **Condition D** | |
| **2AGPE140tipp** | irreversible in forward | blocked |
| **2AGPE141tipp** | irreversible in forward | blocked |
| **2AGPEAT140** | irreversible in forward | blocked |
| **2AGPEAT141** | irreversible in forward | blocked |
| **ACACT6r** | bidirectional | irreversible in forward |
| **AGPAT140** | irreversible in forward | blocked |
| **AGPAT141** | irreversible in forward | blocked |
| **APG3PAT140** | irreversible in forward | blocked |
| **APG3PAT141** | irreversible in forward | blocked |
| **CLPNH140pp** | irreversible in forward | blocked |
| **CLPNH141pp** | irreversible in forward | blocked |
| **CLPNS140pp** | irreversible in forward | blocked |
| **CLPNS141pp** | irreversible in forward | blocked |
| **CTECOAI6** | bidirectional | irreversible in backward |
| **ECOAH6** | bidirectional | irreversible in forward |
| **FACOAL140t2pp** | irreversible in forward | blocked |
| **FACOAL141t2pp** | irreversible in forward | blocked |
| **G3PAT140** | irreversible in forward | blocked |
| **G3PAT141** | irreversible in forward | blocked |
| **HACD6** | bidirectional | irreversible in forward |
| **LPLIPAL1A140pp** | irreversible in forward | blocked |
| **LPLIPAL1A141pp** | irreversible in forward | blocked |
| **LPLIPAL1E140pp** | irreversible in forward | blocked |
| **LPLIPAL1E141pp** | irreversible in forward | blocked |
| **LPLIPAL1G140pp** | irreversible in forward | blocked |
| **LPLIPAL1G141pp** | irreversible in forward | blocked |
| **PA140abcpp** | irreversible in forward | blocked |
| **PA141abcpp** | irreversible in forward | blocked |
| **LPLIPAL2E140** | irreversible in forward | blocked |
| **LPLIPAL2E141** | irreversible in forward | blocked |
| **PE140abcpp** | irreversible in forward | blocked |
| **PE141abcpp** | irreversible in forward | blocked |
| **PG140abcpp** | irreversible in forward | blocked |
| **PG141abcpp** | irreversible in forward | blocked |
| **PGP140abcpp** | irreversible in forward | blocked |
| **PGP141abcpp** | irreversible in forward | blocked |
| **PGPP140** | irreversible in forward | blocked |
| **PGPP140pp** | irreversible in forward | blocked |
| **PGPP141** | irreversible in forward | blocked |
| **PGPP141pp** | irreversible in forward | blocked |
| **PLIPA2A140pp** | irreversible in forward | blocked |
| **PLIPA2A141pp** | irreversible in forward | blocked |
| **PLIPA2E140pp** | irreversible in forward | blocked |
| **PLIPA2E141pp** | irreversible in forward | blocked |
| **PGSA140** | irreversible in forward | blocked |
| **PGSA141** | irreversible in forward | blocked |
| **PLIPA2G140pp** | irreversible in forward | blocked |
| **PLIPA2G141pp** | irreversible in forward | blocked |
| **PLIPA1E140pp** | irreversible in forward | blocked |
| **PLIPA1E141pp** | irreversible in forward | blocked |
| **PSD140** | irreversible in forward | blocked |
| **PSD141** | irreversible in forward | blocked |
| **PSSA140** | irreversible in forward | blocked |
| **PSSA141** | irreversible in forward | blocked |

Table S4 :Changed reactions directionalities after constraining metabolite concentrations in$strain \gamma$ under different conditions.

| **Reaction IDs** | **Unconstrained metabolite concentrations** | **Constrained metabolite concentrations** |
| --- | --- | --- |
|  | **Condition A** | |
| **2AGPE140tipp** | irreversible in forward | blocked |
| **2AGPE141tipp** | irreversible in forward | blocked |
| **2AGPEAT140** | irreversible in forward | blocked |
| **2AGPEAT141** | irreversible in forward | blocked |
| **ACACT6r** | bidirectional | irreversible in forward |
| **AGPAT140** | irreversible in forward | blocked |
| **AGPAT141** | irreversible in forward | blocked |
| **APG3PAT140** | irreversible in forward | blocked |
| **APG3PAT141** | irreversible in forward | blocked |
| **CLPNH140pp** | irreversible in forward | blocked |
| **CLPNH141pp** | irreversible in forward | blocked |
| **CLPNS140pp** | irreversible in forward | blocked |
| **CLPNS141pp** | irreversible in forward | blocked |
| **CTECOAI6** | bidirectional | irreversible in backward |
| **ECOAH6** | bidirectional | irreversible in forward |
| **FACOAL140t2pp** | irreversible in forward | blocked |
| **FACOAL141t2pp** | irreversible in forward | blocked |
| **G3PAT140** | irreversible in forward | blocked |
| **G3PAT141** | irreversible in forward | blocked |
| **HACD6** | bidirectional | irreversible in forward |
| **LPLIPAL1A140pp** | irreversible in forward | blocked |
| **LPLIPAL1A141pp** | irreversible in forward | blocked |
| **LPLIPAL1E140pp** | irreversible in forward | blocked |
| **LPLIPAL1E141pp** | irreversible in forward | blocked |
| **LPLIPAL1G140pp** | irreversible in forward | blocked |
| **LPLIPAL1G141pp** | irreversible in forward | blocked |
| **PA140abcpp** | irreversible in forward | blocked |
| **PA141abcpp** | irreversible in forward | blocked |
| **LPLIPAL2E140** | irreversible in forward | blocked |
| **LPLIPAL2E141** | irreversible in forward | blocked |
| **PE140abcpp** | irreversible in forward | blocked |
| **PE141abcpp** | irreversible in forward | blocked |
| **PG140abcpp** | irreversible in forward | blocked |
| **PG141abcpp** | irreversible in forward | blocked |
| **PGP140abcpp** | irreversible in forward | blocked |
| **PGP141abcpp** | irreversible in forward | blocked |
| **PGPP140** | irreversible in forward | blocked |
| **PGPP140pp** | irreversible in forward | blocked |
| **PGPP141** | irreversible in forward | blocked |
| **PGPP141pp** | irreversible in forward | blocked |
| **PLIPA2A140pp** | irreversible in forward | blocked |
| **PLIPA2A141pp** | irreversible in forward | blocked |
| **PLIPA2E140pp** | irreversible in forward | blocked |
| **PLIPA2E141pp** | irreversible in forward | blocked |
| **PGSA140** | irreversible in forward | blocked |
| **PGSA141** | irreversible in forward | blocked |
| **PLIPA2G140pp** | irreversible in forward | blocked |
| **PLIPA2G141pp** | irreversible in forward | blocked |
| **PLIPA1E140pp** | irreversible in forward | blocked |
| **PLIPA1E141pp** | irreversible in forward | blocked |
| **PSD140** | irreversible in forward | blocked |
| **PSD141** | irreversible in forward | blocked |
| **PSSA140** | irreversible in forward | blocked |
| **PSSA141** | irreversible in forward | blocked |
|  | **Condition B** | |
| **ACACT6r** | bidirectional | irreversible in backward |
| **ACACT7r** | irreversible in forward | blocked |
| **ACACT8r** | irreversible in backward | blocked |
| **ACOAD7f** | irreversible in backward | blocked |
| **CTECOAI6** | bidirectional | irreversible in forward |
| **CTECOAI8** | irreversible in backward | blocked |
| **ECOAH6** | bidirectional | irreversible in backward |
| **ECOAH7** | irreversible in forward | blocked |
| **ECOAH8** | irreversible in forward | blocked |
| **FACOAE160** | irreversible in forward | blocked |
| **FACOAE181** | irreversible in forward | blocked |
| **HACD6** | bidirectional | irreversible in backward |
| **HACD7** | irreversible in forward | blocked |
| **HACD8** | irreversible in forward | blocked |
| **OCTNLL** | irreversible in forward | blocked |
|  | **Condition C** | |
| **ACACT6r** | bidirectional | irreversible in backward |
| **ACACT7r** | irreversible in forward | blocked |
| **ACACT8r** | irreversible in backward | blocked |
| **ACOAD7f** | irreversible in backward | blocked |
| **CTECOAI6** | bidirectional | irreversible in forward |
| **CTECOAI8** | irreversible in backward | blocked |
| **ECOAH6** | bidirectional | irreversible in backward |
| **ECOAH7** | irreversible in forward | blocked |
| **ECOAH8** | irreversible in forward | blocked |
| **FACOAE160** | irreversible in forward | blocked |
| **FACOAE181** | irreversible in forward | blocked |
| **HACD6** | bidirectional | irreversible in backward |
| **HACD7** | irreversible in forward | blocked |
| **HACD8** | irreversible in forward | blocked |
|  | **Condition D** | |
| **2AGPE140tipp** | irreversible in forward | blocked |
| **2AGPE141tipp** | irreversible in forward | blocked |
| **2AGPEAT140** | irreversible in forward | blocked |
| **2AGPEAT141** | irreversible in forward | blocked |
| **ACACT6r** | bidirectional | irreversible in forward |
| **AGPAT140** | irreversible in forward | blocked |
| **AGPAT141** | irreversible in forward | blocked |
| **APG3PAT140** | irreversible in forward | blocked |
| **APG3PAT141** | irreversible in forward | blocked |
| **CLPNH140pp** | irreversible in forward | blocked |
| **CLPNH141pp** | irreversible in forward | blocked |
| **CLPNS140pp** | irreversible in forward | blocked |
| **CLPNS141pp** | irreversible in forward | blocked |
| **CTECOAI6** | bidirectional | irreversible in backward |
| **ECOAH6** | bidirectional | irreversible in forward |
| **FACOAL140t2pp** | irreversible in forward | blocked |
| **FACOAL141t2pp** | irreversible in forward | blocked |
| **G3PAT140** | irreversible in forward | blocked |
| **G3PAT141** | irreversible in forward | blocked |
| **HACD6** | bidirectional | irreversible in forward |
| **LPLIPAL1A140pp** | irreversible in forward | blocked |
| **LPLIPAL1A141pp** | irreversible in forward | blocked |
| **LPLIPAL1E140pp** | irreversible in forward | blocked |
| **LPLIPAL1E141pp** | irreversible in forward | blocked |
| **LPLIPAL1G140pp** | irreversible in forward | blocked |
| **LPLIPAL1G141pp** | irreversible in forward | blocked |
| **PA140abcpp** | irreversible in forward | blocked |
| **PA141abcpp** | irreversible in forward | blocked |
| **LPLIPAL2E140** | irreversible in forward | blocked |
| **LPLIPAL2E141** | irreversible in forward | blocked |
| **OCTNLL** | irreversible in forward | blocked |
| **PE140abcpp** | irreversible in forward | blocked |
| **PE141abcpp** | irreversible in forward | blocked |
| **PG140abcpp** | irreversible in forward | blocked |
| **PG141abcpp** | irreversible in forward | blocked |
| **PGP140abcpp** | irreversible in forward | blocked |
| **PGP141abcpp** | irreversible in forward | blocked |
| **PGPP140** | irreversible in forward | blocked |
| **PGPP140pp** | irreversible in forward | blocked |
| **PGPP141** | irreversible in forward | blocked |
| **PGPP141pp** | irreversible in forward | blocked |
| **PLIPA2A140pp** | irreversible in forward | blocked |
| **PLIPA2A141pp** | irreversible in forward | blocked |
| **PLIPA2E140pp** | irreversible in forward | blocked |
| **PLIPA2E141pp** | irreversible in forward | blocked |
| **PGSA140** | irreversible in forward | blocked |
| **PGSA141** | irreversible in forward | blocked |
| **PLIPA2G140pp** | irreversible in forward | blocked |
| **PLIPA2G141pp** | irreversible in forward | blocked |
| **PLIPA1E140pp** | irreversible in forward | blocked |
| **PLIPA1E141pp** | irreversible in forward | blocked |
| **PSD140** | irreversible in forward | blocked |
| **PSD141** | irreversible in forward | blocked |
| **PSSA140** | irreversible in forward | blocked |
| **PSSA141** | irreversible in forward | blocked |
